# CytoBatchNorm: an R package with graphical interface for batch effects correction of cytometry data

**DOI:** 10.1101/2024.05.29.596492

**Authors:** Samuel Granjeaud, Naoill Abdellaoui, Anne-Sophie Chrétien, Eloise Woitrain, Laurent Pineau, Sandro Ninni, Alexandre Harari, Marion Arnaud, David Montaigne, Bart Staels, David Dombrowicz, Olivier Molendi-Coste

## Abstract

Innovation in cytometry propelled it to an almost “omic” dimension technique during the last decade. The application fields concomitantly enlarged, resulting in generation of high-dimensional high-content data sets which have to be adequately designed, handled and analyzed. Experimental solutions and detailed data processing pipelines were developed to reduce both the staining conditions variability between samples and the number of tubes to handle. However, an unavoidable variability appears between samples, barcodes, series and instruments (in multicenter studies) contributing to “batch effects” that must be properly controlled. Computer aid to this aim is necessary, and several methods have been published so far, but configuring and carrying out batch normalization remains unintuitive for scientists with “pure” academic backgrounds in biology. To address this challenge, we developed an R package called CytoBatchNorm that offers an intuitive and user-friendly graphical interface. Although the processing is based on the script by Schuyler et al., the graphical interface revolutionizes its use. CytoBatchNorm enables users to define a specific correction for each marker in a single run. It provides a graph that guides you through quickly setting the correction for each marker. It allows corrections to be previewed and inter-marker effects to be checked as the settings are made. CytoBatchNorm will help the cytometry community to adequately scale data between batches, reliably reducing batch effects and improving subsequent dimension reduction and clustering.

**VISUAL ABSTRACT:** 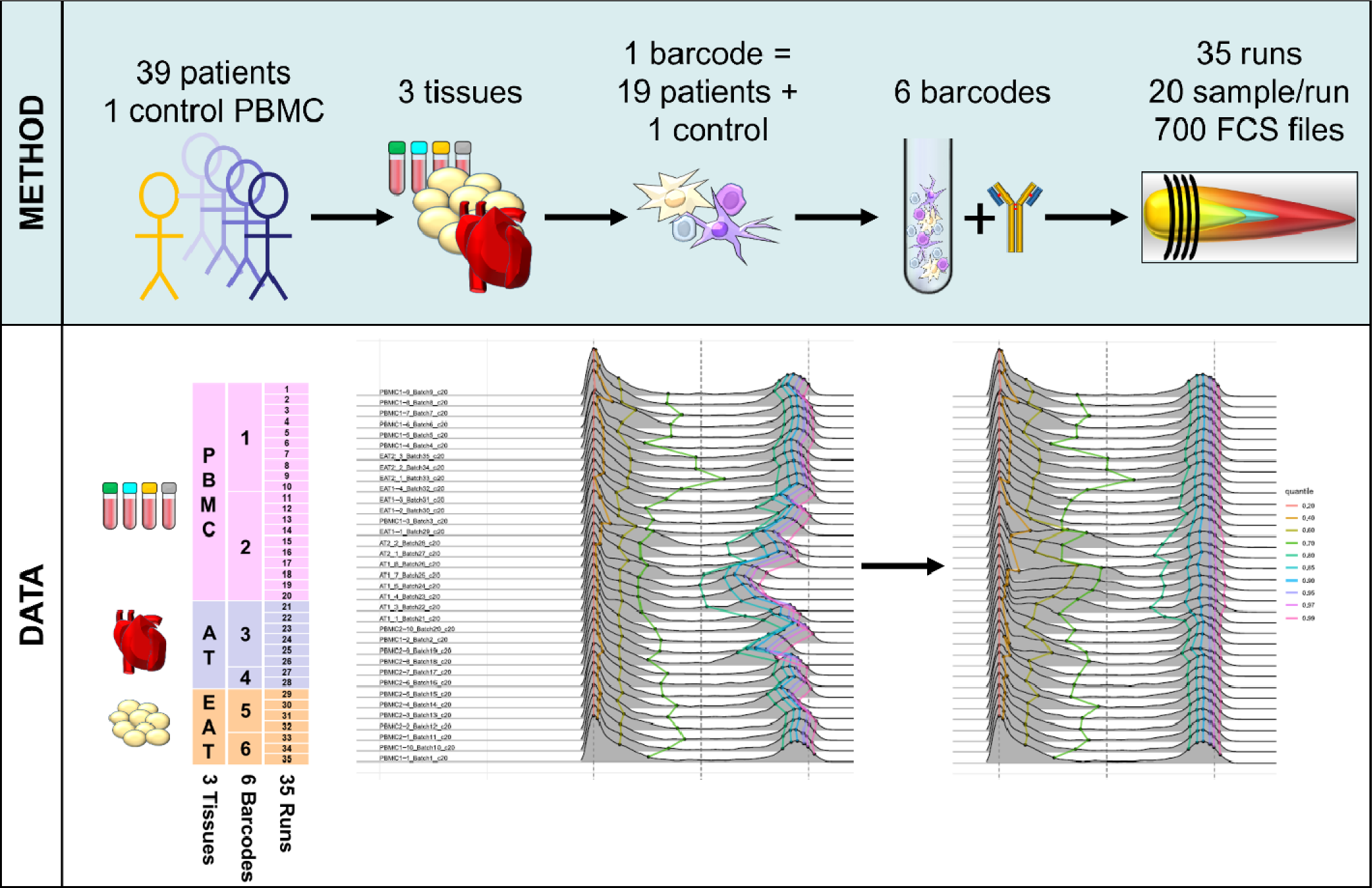

## INTRODUCTION

Innovation in cytometry propelled it to an almost “omic” dimension technique during the last decade, with the rising of both cytometry by time-of-flight (CyTOF, or mass cytometry) and spectral cytometry, allowing elaboration of panels up to 30-50 parameters analyzed at the single cell level[1]. The application fields concomitantly enlarged, with data sets that include more and more (i) samples - specifically in human clinical studies, (ii) tissues from a single donor, and (iii) stimulated conditions for a single sample, all leading to an increase in the number of “tubes” composing an experiment. This results in generation of high-dimensional high-content data which have to be adequately designed, handled and analyzed[2].

Experimental solutions and detailed samples processing pipelines were developed to reduce both the staining conditions variability between tubes[3] and the number of tubes to handle/acquire, the later mainly consisting in barcoding of samples[4,5]. However, barcoding has its intrinsic limits in the number of samples that can be barcoded together, and acquisition of a single barcode (composed of tens of millions of cells) is very long and usually achieved in many runs, notably in mass cytometry in regard with its limited acquisition rate speed. Thus, large datasets are usually obtained during acquisitions spanning days or weeks. Despite the maximal attention paid by experimenters in standardizing their protocols, sequential manipulation and instrument instability over time[3] introduce variability between samples, barcodes and runs. The raise of multi-center studies also introduces an instrument-dependent effect that needs to be corrected to enhance data consistency[6]. All these sources of variations in results baseline participate to the commonly used “batch effects” denomination and have to be controlled properly[7–11].

Batch effects can be addressed at two levels of the experimental protocol. The instrument-related batch effect is mainly dependent on tuning and quality control (QC) accuracy, specific instrument sensitivity and sensitivity variations along acquisitions. Normalization methods based on calibration beads were developed for both flow[12,13] and mass cytometry[5] to counteract this variability level, and EQbeads (Standard BioTools Inc, San Francisco, CA, US) were structurally included in mass cytometers design. However, these methods have their own limitations as illustrated below and by others[3,14]. Based on external controls (beads), they only aim to standardize the instrument. The experimental-related batch effects basically regroup all parameters of an experiment that can influence staining and detection efficiencies of samples, in addition to the instrument used for acquisition.

To evaluate signal stability along batches/days, solutions have been developed, such as the inclusion of a “control” aliquot (i.e. a standard sample) in all batches. This design served as a substrate for the development of algorithms to correct batch effects[9]. Methods and software packages published so far[15–23] circumvent batch variations in channels intensity using either an “all-events” or a “pre-gated population-specific” adjustment. Whatever the package/method considered, configuration as well as application of the normalization process across batches remains either obscure or unintuitive for scientists with “pure” academic background in biology (Table 1), which usually and historically lack training in computer science and mathematics.

**Table 1:** Main published packages for batch effect correction. 1: as no batch is defined, all samples are aligned; when batch information is available, the transform defined within each batch is applied to every sample of the batch, which removes differences between batches but keeps differences within each batch.

| Method | Based on | Define batches | Includes a clustering | Requires a standard sample | Align | Parameter to be chosen | Visual interface | Normalization preview |
| --- | --- | --- | --- | --- | --- | --- | --- | --- |
| iMuBAC | harmony | ✓ | ✓ |  |  | auto | No | No |
| gaussNorm, fdaNorm | - | ✗ <sup>1</sup> |  |  | detected peaks | auto | No | No |
| cyCombine | Combat | ✓ | ✓ |  |  | auto | No | No |
| CytoNorm | - | ✓ | ✓ | ✓ | 101 quantiles | auto | No | No |
| cytofRUV | RUV | ✓ |  |  |  | depth | No | No |
| CytofBatchAdjust | - | ✓ |  | ✓ | 1 percentile | percentile | No | No |
| CytoBatchNorm | CytofBatchAdjust | ✓ |  | ✓ | 1 (or 2) percentiles | percentile | Yes | Yes |

The CytofBatchAdjust R script released by Schuyler et al. in 2019[19] smartly proposed an “all-events” adjustment of peaks intensities for each channel of the control tube included in each batch to the peaks intensities of the control tube of a referent batch. This adjustment is performed channel by channel, independently of each other. It can be achieved with different options, scaling on quantiles or on a specified percentile, considering or not zero values and with or without arcsinh transformation of data. As depicted by the authors, the “quantile” adjustment is not adequate for normalization of cytometry data, because it sometimes creates artefacts that can only be identified on a bi-parametric plot. Indeed, when the distribution of events in the control sample differs even slightly between the reference batch and the other batches (e.g. the proportion of events in the positive and negative peaks), the “quantile” adjustment shifts events from one peak to the other. By contrast, the “percentile” method allows the user to choose a single percentile value (from 1 to 99) which identifies an intensity in each batch and linearly scales each batch in order to align those intensities to the intensity of the reference batch. Determining the best percentile for a given channel has so far been tricky and empirical, mainly because users lack an indicative graph. If the percentile is chosen in a region where the distribution of events varies between the control tubes of different batches, the channel scaling will be aberrant in some batches, introducing computational biases in the downstream data analysis, which would only be identified after the correction has been applied.

To provide a comprehensive, transparent and interactive pipeline to reliably overcome the different levels of batch variability, we developed the CytoBatchNorm R package, which is adapted from CytofBatchAdjust code and methodology[19]. We illustrate its use with an experiment of 700 FSC files from 35 batches including 3 human tissues and benchmark it on two different data sets versus the two most-commonly used packages: CytofBatchAdjust[19] and CytoNorm[18]. This package is intended to be “ready-to-use” for experimenters without any R coding skills. We improved the CytofBatchAdjust R script by providing (i) an intuitive user-friendly interface, (ii) an interactive graphic for each marker to help selecting the best percentile relative to the distribution of intensity across the control samples in each batch before proceeding with normalization, (iii) a comprehensive table to specify the percentile channel-by-channel, (iv) the ability to perform a bi-percentile adjustment, (v) dot-plots and output graphs to control the batch adjustment accuracy, (vi) code for the Windows system.

## METHODS

### Dataset 1

Datasets structures are summarized in Table 2

**Table 2:**
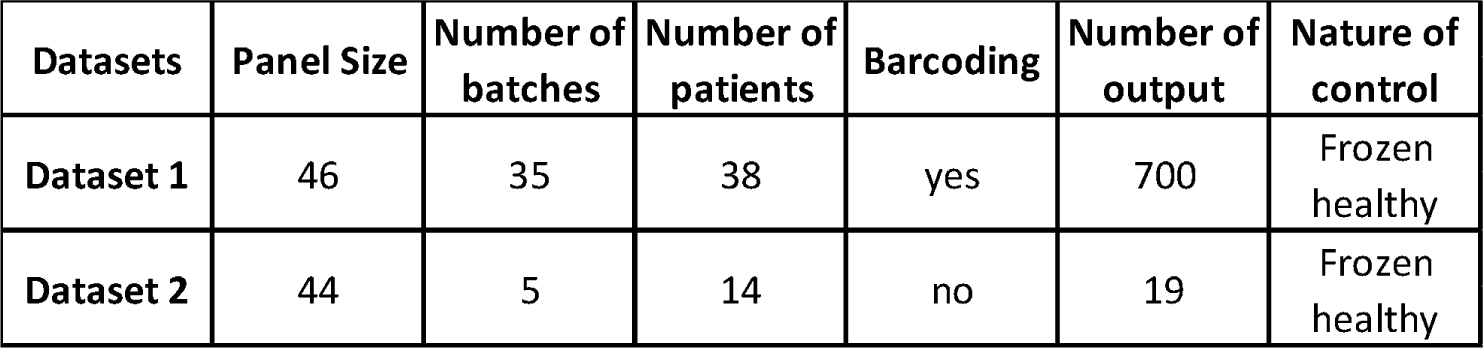
Datasets structure.

#### Sample collection

Atrial Myocardial Tissue (AT) obtained from the right atrial appendage before aortic cross-clamping and cardioplegia, Epicardic Adipose Tissue (EAT) and PBMC from 38 patients undergoing a Surgical Aortic Valve Replacement (SAVR) (POMI-AFclinical study NCT#03376165; PI: D Montaigne) as well as PBMC from a single healthy donor were analyzed by mass cytometry. Written informed consent was obtained from all patients before inclusion. Non-parenchymal cells fraction from AT and EAT were prepared from tissues immediately cleaned, minced and digested at 37°C for 45 min in 0.1% collagenase I and PBMC were obtained as previously described[24].

#### Staining protocol

Detailed antibody panel and staining protocol were described previously[24]. Briefly, thawed PBMCs (3x10^6^ cells) and non-parenchymal cells from AT and EAT (40x10^3^ to 1.4.10^6^ cells) were sequentially stained for viability, for extracellular targets sensitive to fixatives, barcoded, stained for extracellular markers not sensitive to fixatives, for intracellular targets and for DNA as summarized in Table 3. Six independent pools of 19 experimental samples (2 pools of PBMC, 2 pools of AT, 2 pools of EAT) added with one aliquot of the control PBMC sample from a single healthy donor were processed independently for barcoding (each pool for a total of 20 samples), as summarized in Graphical Abstract.

**Table 3:**
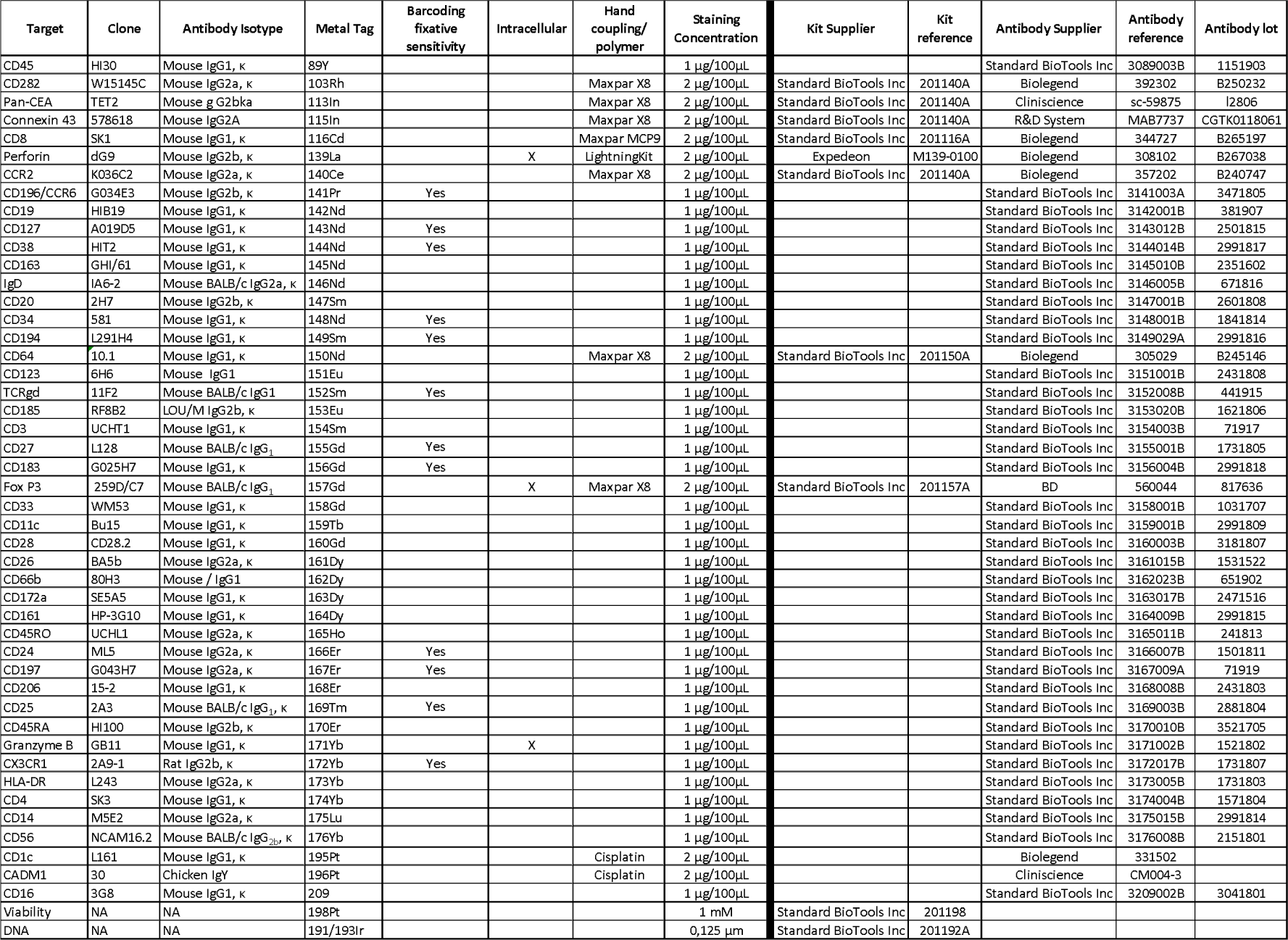
Antibody panel from Dataset 1.

#### Acquisition

Cells were resuspended in Maxpar Water at 5x10^5^ cells/mL with 1:10 volume of Four-Element Calibration Beads (Standard BioTools Inc, San Francisco, CA, US) and analyzed on an Helios instrument (Standard BioTools Inc, San Francisco, CA, US). Runs were realized for a maximum of 150 minutes at a maximum rate of 500 events/sec, and a quick tuning was performed between each run.

#### Datasets 2

Datasets structures are summarized in Table 2

#### Sample collection

*In vitro*-expanded tumor-infiltrating lymphocytes (TILs) were generated from tumor samples of 14 cancer patients, using an improved TIL culture method [25]. PBMCs from a single healthy donor were used as control sample. Samples were collected and biobanked from patients enrolled under protocols approved by the Lausanne university hospital (CHUV), Switzerland. Patients and healthy donors’ recruitment, study procedures, and blood withdrawal were approved by regulatory authorities and all patients signed written informed consents.

#### Staining protocol

Frozen TILs (1x10^6^ to 5.10^6^ cells) and PBMCs (3x10^6^ cells) were thawed, rested overnight, and labelled with 44 metal-coupled antibodies (Standard BioTools Inc, San Francisco, CA, US & in house). Cells were first stained for viability then stained for extracellular targets (Standard BioTools 400276 Protocol).

Cells were then fixed using Cytofix fixation buffer (BD Biosciences) and permeabilized using Phosflow Perm Buffer III solution (BD Biosciences). Next, intracellular staining was performed (antibody incubation: 30 min at RT). For DNA staining, cells were next incubated overnight at 4°C with cell intercalation solution following the manufacturer’s protocol (Standard BioTools 400276 Protocol).

#### Acquisition

Cells were finally washed and resuspended in a Maxpar Cell Acquisition Solution containing EQ Four Element Calibration Beads (Standard BioTools Inc, San Francisco, CA, US) at a cell concentration of 10^6^ cells/mL, immediately prior to CyTOF data acquisition. Five runs at different days were realized using an Helios Mass Cytometer (Standard BioTools Inc, San Francisco, CA, US) at a maximum rate of 500 events/sec.

### Data processing

Raw mass cytometry data (Datasets 1 and 2) were first normalized with the calibration EQ-bead passport pre-loaded in the CyTOF Software version 7 (Standard BioTools 400276 Protocol) and then debarcoded following the manufacturer’s instructions (Dataset 1). Data pre-processing was performed using Cytobank (Beckman Coulter, Indianapolis, IN, USA) for Dataset 1 or FlowJo 10 (Becton Dickinson, Franklin Lakes, NJ, US) for Dataset 2.

### Batch effect normalization

#### Dataset 1

The 6 barcoded pools of samples all included an aliquot of a control PBMC sample from a single healthy donor (C20). Those 6 barcoded pools were acquired on an Helios instrument through a total of 35 different runs dispatched as follows: PBMC pool 1 – 10 runs; PBMC pool 2 – 10 runs; AT pool 1 – 6 runs; AT pool 2 – 2 runs; EAT pool 1 – 4 runs; EAT pool 2 – 3 runs (Graphical Abstract). Those 35 runs were thereafter considered as 35 independent batches in order to compensate for possible signal drift over duration of acquisition of a single barcoded pool.

The C20 included in each barcoded run was gated on nucleated - single - biological - non beads - CD45^+^ live events (Supplemental Figure 1) and used as an anchor for application of the modified CytofBatchAdjust R code published by Schuyler et al.[19] as exposed in the RESULTS section. Batch-adjusted FCS files from each single sample were concatenated to reconstitute original samples.

#### Dataset 2

Five batches of samples, all including an aliquot of a control PBMC sample from a single healthy donor, were acquired on an Helios instrument over a total of 5 different runs (1 run per batch). The PBMC from the healthy donor included in each batch was gated on nucleated - single - biological - non beads - CD45^+^ live events (Supplemental Figure 1) and used as an anchor for application of the modified CytofBatchAdjust R code published by Schuyler et al.[19]. Dataset 2 served for benchmarking against CytoNorm[18], which was performed as a plugin supplied by FlowJo software (Becton Dickinson, Franklin Lakes, NJ, US) following the provider’s instructions.

### Spillover compensation

#### Dataset 1

For compensation matrix calculation, Comp Beads (Becton Dickinson, Franklin Lakes, NJ, US) were single stained in CSB with 1µg of each antibody for 30 minutes at RT. Exception was made for the anti-CADM1 antibody labelled with 196Pt which is a chicken IgY, that is not captured by Comp Beads, which was replaced by an anti-CD8 (clone SK1, Mouse IgG1, κ) conjugated with the same batch of 196Pt. After two washes with CSB and two wash with Marxpar Water, beads were mixed and acquired as a single tube at a maximum rate of 500 events/sec. The FCS file from mixed single stained beads was imputed in CATALYST R[26] using the NNLS method. The output compensation matrix (Figure 6 C) was applied to all the files and compensated FCS files were edited and processed to data analysis.

### Data analysis

Debarcoded - batch normalized – concatenated - compensated FCS files were gated on nucleated - single - biological - non beads - CD45^+^ live events (Supplemental figure 1) and processed for phenotype analysis using R 4.0.0 and a modified version of the Cytofkit package[27] (http://github.com/i-cyto/cytofkitlab), including UMAP computation using the uwot package. Dimension reduction was performed using t-SNE or UMAP algorithms. Multi-dimensional scaling was performed using the CytoMDS R package[28]. Illustrations were edited using the Cytofkit ShinyAPP browser[27] and Cytobank (Beckman Coulter, Indianapolis, IN, USA).

### Script

The package as well as recommendations and commands are described here: https://github.com/i-cyto/cytoBatchNorm. Installing and launching of the package are simply done using the commands below in R or R Studio.

Donwlaod/installation:

devtools::install_github(“i-cyto/cytoBatchNorm”)

Launching:

library(cytoBatchNorm)

cytoBatchNormGUI()

## RESULTS and DISCUSSION

Computer-assisted treatment of data, whatever the kind of treatment, has to be finely tuned and controlled carefully to avoid computational artefacts as well as data distortions that could deteriorate data consistency and induce uncontrolled bias in the downstream analysis pipeline. Considering batch effects correction, this burden should be taken in consideration cautiously especially considering the complexity of the dataset (number of batches, samples, markers), as both the risk of such data distortion and the possibility that the experimenter misses it during visual control of corrected data increase with the dimensionality of the dataset.

As a rational and accessible solution, we developed an intuitive, user-friendly tool based on R Shiny package that does not require any practice in R language or programming skills. This tool can easily be installed and launched using simple commands referenced in the “script” section. The Web interface allows intuitive, point-and-click navigation through the different steps of the normalization pipeline: selection of the dataset (Figure 1A), identification of the different batches/control tubes and of the referent control tube (Figure 1B), selection of the channels and tuning of the batch correction process (Figure 1C) as well as launching the correction.

**Figure 1:**
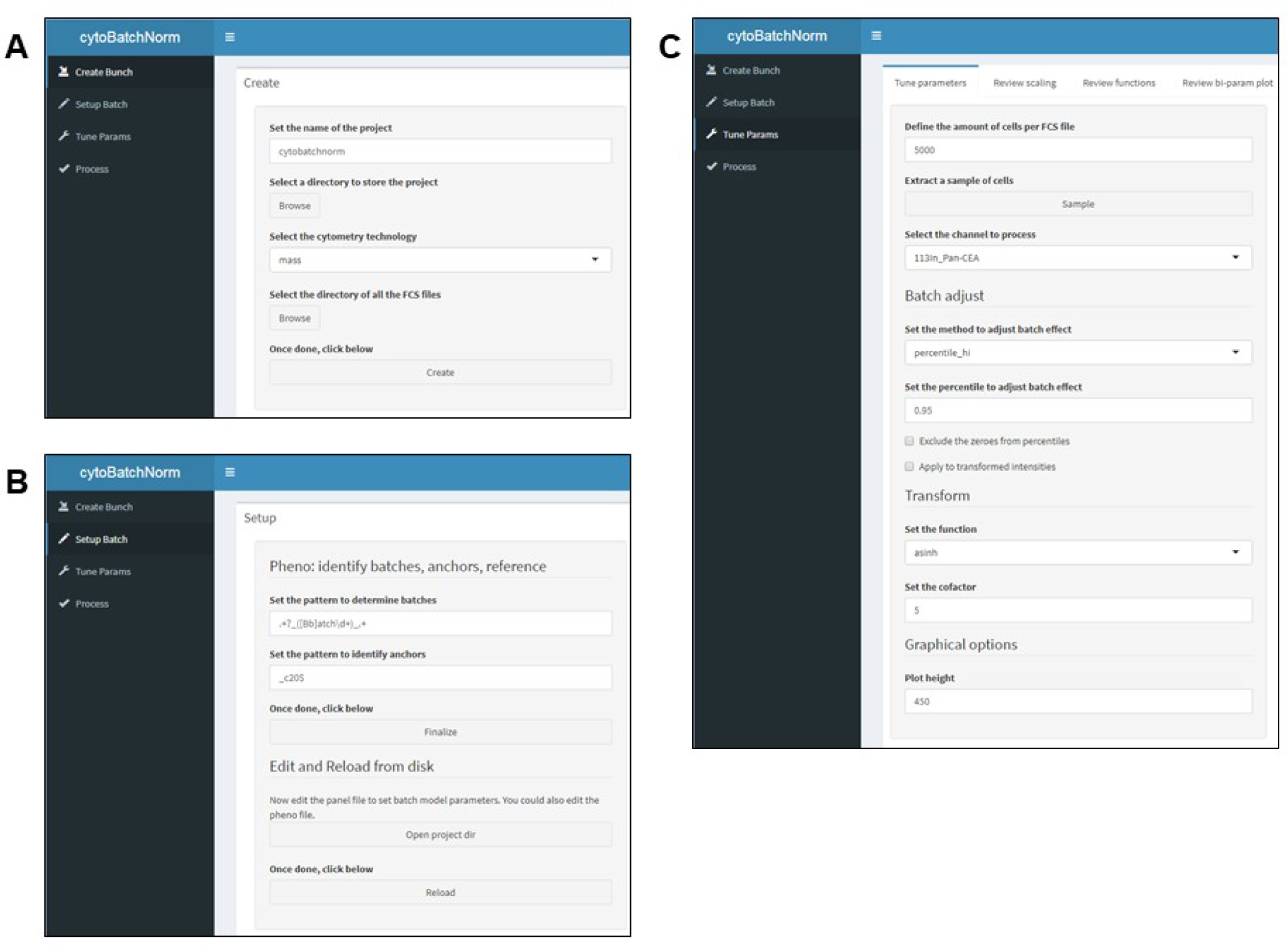
Sequential operation of the batch effect normalization interface. A: Bunch creation menu. B: Batch setup menu (detailed in figure 2). C: Tuning parameters menu (detailed in figure 3 and 4).

### “Create Bunch” menu

The “Create Bunch” menu (Figure 1A) asks for the name of the current experiment to be created, storage directory, cytometry technology as well as the directory containing the FCS files to adjust. Finalization is done by clicking on the “Create” button.

### “Setup Batch” menu

The “Setup Batch” menu (Figure 1B) allows to define keywords for automatic identification of batches and control tubes (termed “anchor”, “C20” in our example) for each batch inside the FCS files directory. The “finalize” button launches the identification process and creates two tables (.xlsx) “pheno” and “panel” which can be accessed using the “open project dir” button (Figure 1B and Figure 2A-2). Those tables can be edited and saved using appropriate spreadsheets software, and then updated in the interface by clicking the “reload” button (Figure 2A-2). The “pheno” table (Figure 2A-3) lists the files identified as control tubes (column “sample_ID_is_ref” = “Y”) and the reference batch control tube (column “sample_ID_is_ref” = “Y” and column “batch_is_ref” = “Y”). The choice of the reference batch is decided by the user, setting only one “Y” in the “batch_is_ref” column onto the proper row/file in the .xlsx “pheno” table, saving it, and then clicking on the “reload” button. The “panel” table (Figure 2B) summarizes the attributes of the FCS files loaded as well as the percentile value that will be used for each channel’s adjustment (by default 0.95), and can be edited the same way as the “pheno” table, if necessary, saved and reloaded similarly.

**Figure 2:**
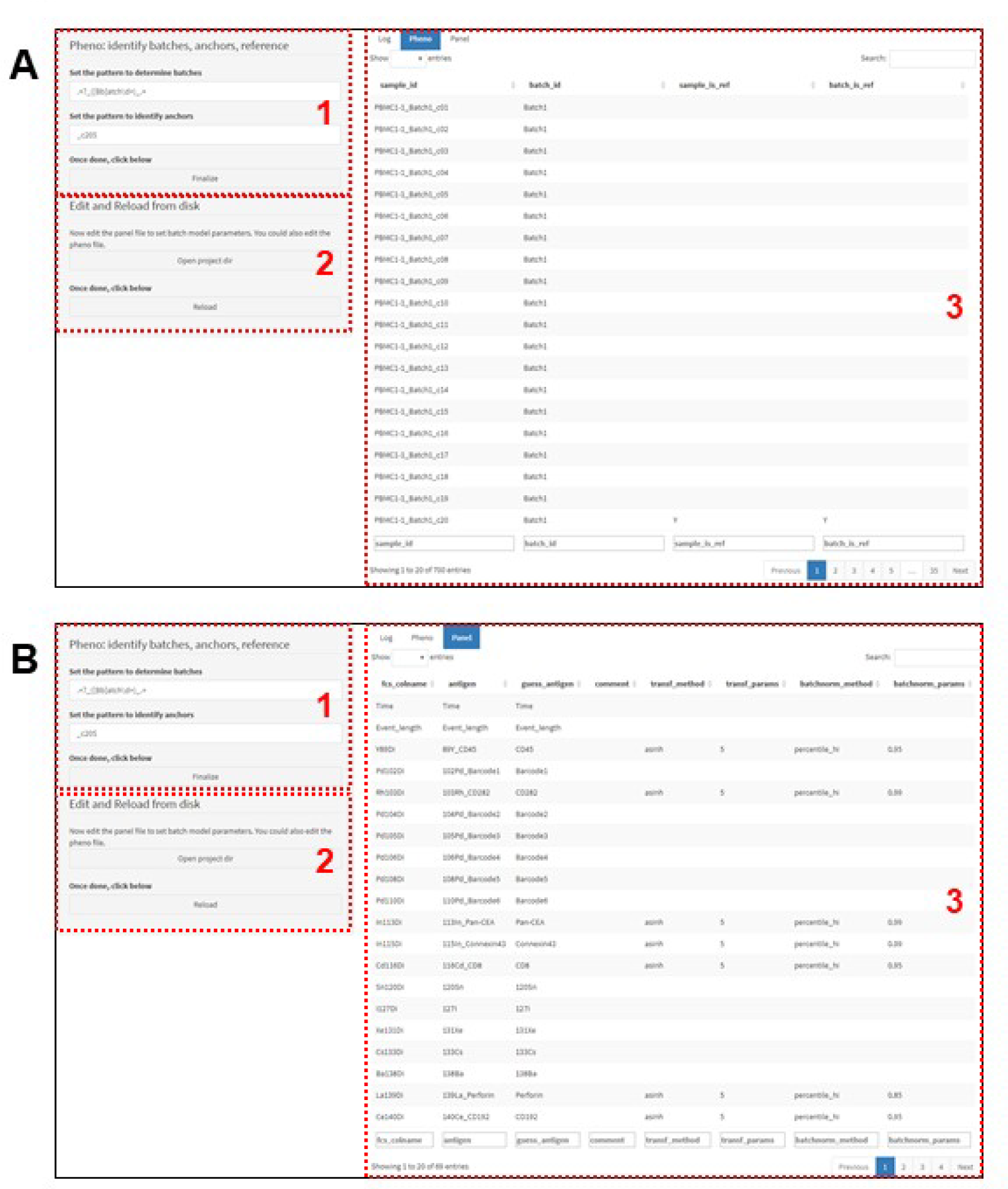
Batch setup menu. A: Tab for setup of the phenotype of the data. B: Tab for setup of the panel of the data. First step (A1, B1) is the identification of the batches and the referent control tubes of each batch using keywords. We strongly recommend to clearly standardize keywords in files nomenclature, i.e. « Batch » and « C20 » respectively in the example displayed. Identification is launched by clicking on the « finalize » button. Users can then check if this identification of batches and referent control tubes is accurate in the pheno tab (A3) and if the indexing of the panel is adequate (B3). Those tables can be modified on spreadshits using the .xlsx files accessible in the project directory, and reloaded using the « reload » button (A2, B2).

**Figure 3:**
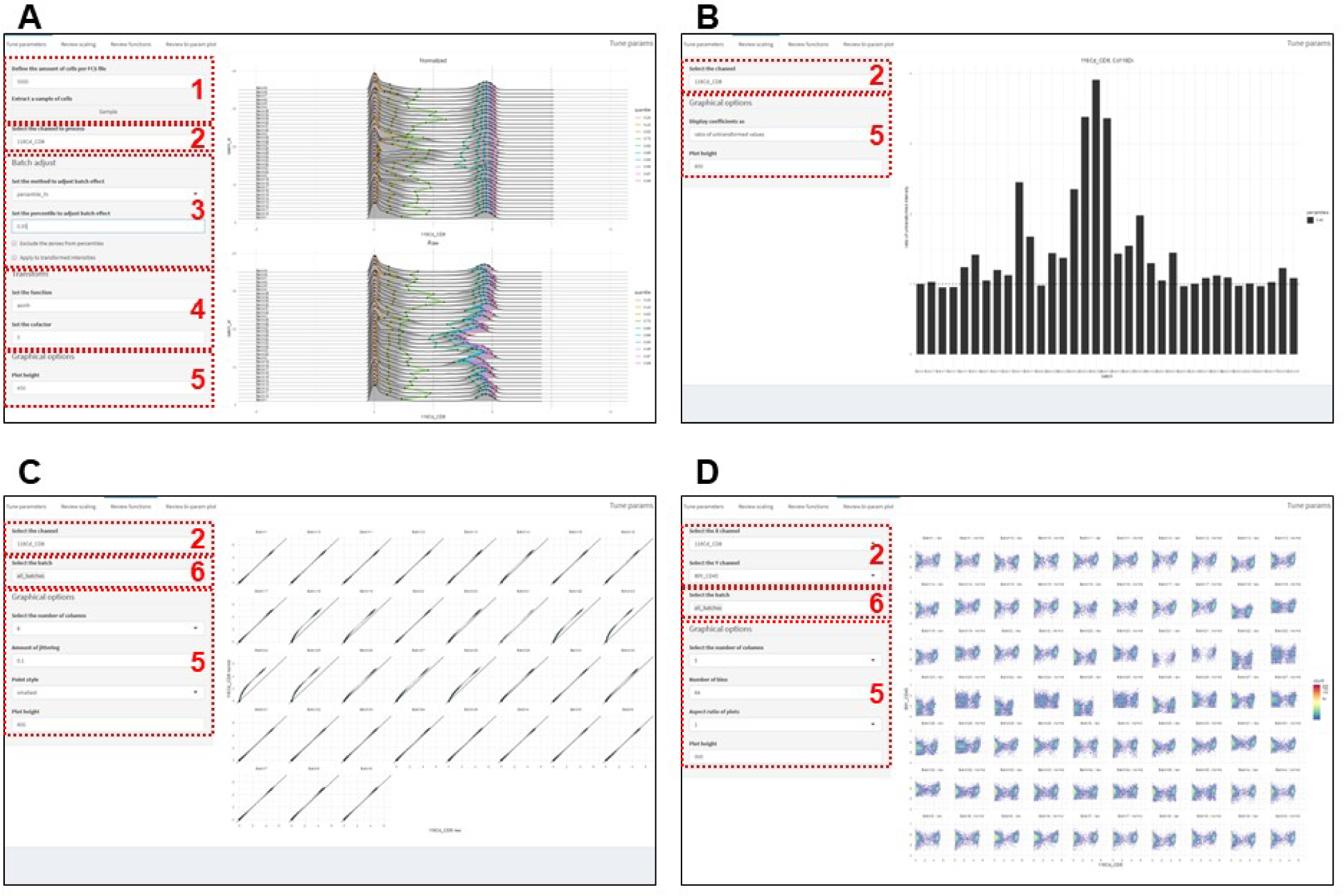
Detailed “Tune parameters” tabs. A: “Tune parameters” main tab. B: “Review scaling” tab. C: “Review functions” tab. D: “Review bi-parameters plot” tab. Tools for parameters tuning include a sampling function (A1), a channels selection function (A2, B2, C2, D2), batch adjustment functions (A3), transform functions (A4), graphical options (A5, B5, C5, D5) and a batch selection function (C6, D6).

### “Tune Params” menu

A major improvement in CytoBatchNorm interface is the addition of an easy “step-by-step” process to determine which percentile will be the most suitable and accurate to scale the whole experiment for each single channel independently. The “Tune parameters” step (Figure 1C and Figure 3) offers a first tab which presents the histograms of all control tubes from all batches for the selected channel in a ridgeline plot and superimposes a line through histograms for each percentile of the default percentile set as illustrated in Figure 3A (0.20, 0.40, 0.60, 0.70, 0.80, 0.90, 0.95, 0.97, 0.99). First, a sampling of files has to be achieved using the “sample” button (Figure 3A-1). Then, users can preview the effect of normalization with different percentiles (Figure 3A-3) and different transformations (Figure 3A-4) in the adjusted and raw intensity graphs. To refine and validate the previewed batch effect correction, we also developed a set of control plots presenting and comparing multiple batches at once. The “Review scaling” tab (Figure 3B) displays the scaling factors that would be applied with the selected parameters. The “Review functions” tab (Figure 3C) shows the raw vs pre-normalized events for the active channel which allows a direct control of the linearity of the normalization. The “Review bi-parameters plot” tab (Figure 3D) allows to build a dot-plots of the active channel vs another channel and to check whether an artefact would be induced by the normalization. These four tabs allow an enlightened determination of which percentile value is the most suitable to scale a given channel, considering the distribution of events along the range, some possible variations in this distribution across batches and notably questioning the relevance of choosing a percentile value in the positive or negative peak or in the positive “queue” if so.

**Figure 4:**
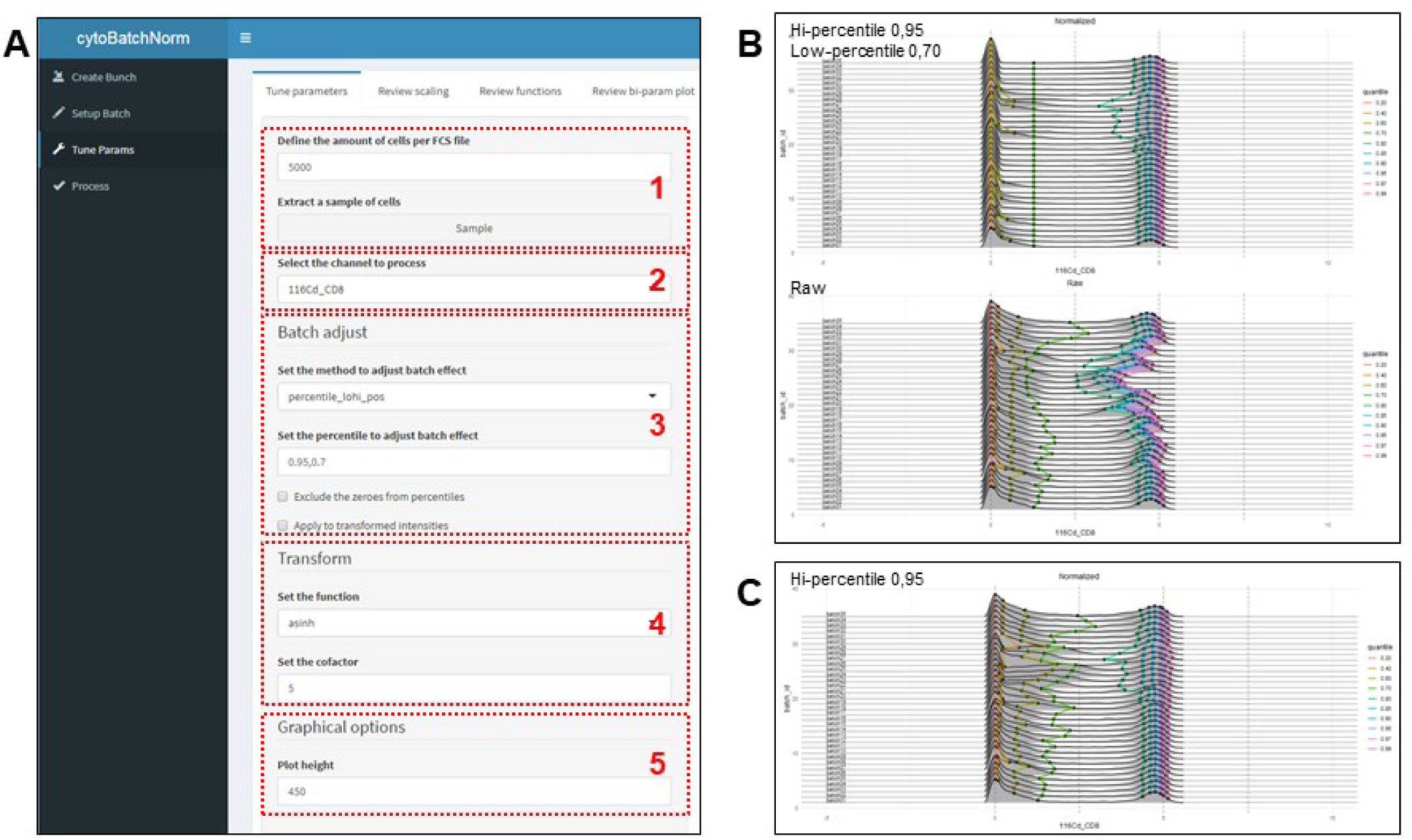
Bi-percentile adjustment. The Batch adjust function in the Tune parameters menu (A-3) allows to choose a “percentile_lohi” option to define two values of percentiles, separated by a comma, which will serve for adjustment of the selected channel (A-2). Predictive modifications of the selected channel are displayed on the right window (B). When compared to a single “percentile_hi” method (C), repartition of the “low” events along the scale range is clearly homogenized, with a highly significant risk to loose linearity of the adjustment within batches. Tools for parameters tuning include sampling functions (1), a channels selection function (2), batch adjustment functions (3), transform functions (4), graphical options (5).

In mass cytometry, zero-values can represent a large part of events depending on the couple marker/metal-tag, which raises questions. Excluding them may greatly change the distribution of percentiles along some channels. Beyond the “zero values” lays the consideration of the negative peak for a given channel, including in the case of unimodal distribution of the events. Indeed, correcting batch effect on the positive peak of a given channel (which is intuitive, as for CD8 in our illustration, Figure 3A) will rescale the entire channel in order to align the positive peak of the batch control tubes to the referent control, shifting values up or down along the whole scale. Depending on the amplitude of the scaling required, it is possible for almost negative events to be moved away from the commonly considered “negative zone” of a channel. To enable users to control this potential adverse effect of a single-percentile adjustment, we implemented a bi-percentile adjustment function, with the idea of defining one percentile for the alignment of the positive peak and another percentile for the alignment of the negative peak. As illustrated in Figure 4, this is simply performed by choosing the “percentile_lohi” option in the “method” menu and entering two percentile values separated with a comma in the “percentile” window (Figure 4A-3). As for the “percentile_hi” option, pre-diagnostic graphs are automatically displayed (Figure 4B). When compared to a single 0.95 percentile adjustment on the positive peak for the CD8 channel of our dataset (Figure 4C), bi-percentile adjustment on 0.95 (positive peak) and 0.70 (light green line in the queue of the negative events) clearly homogenizes the repartition of the events along the scale range, with a highly significant risk to deform data that has to be considered. Indeed, the bi-percentile method is no more linear along the whole scale range, with the same burdens as described above about the quantile method. This “percentile_lohi” method is also provided with the possibility to limit negative adjustment of data to zero (avoid extra-negative values) for mass cytometry, named “percentile_lohi_pos”.

The individual percentiles chosen for each channel in the “Tune Params” menu are retained in the interface and will be applied when clicking on the “preview” button from the “Process menu” (see below). This represents a major improvement, as compared to the original package in which users had to realize multiple successive runs, one for each given percentile value, with specification of which channels had to be adjusted for each specific run. Those channel-specific percentiles can also be easily specified manually in the “Panel” table stored in the project directory (Figure 1A). Doing so, the “Panel” table has to be saved, closed and reloaded manually using the “reload” button (Figure 2A-2 and 2B-2) before going to the “Process” menu (Figure 5).

**Figure 5:**
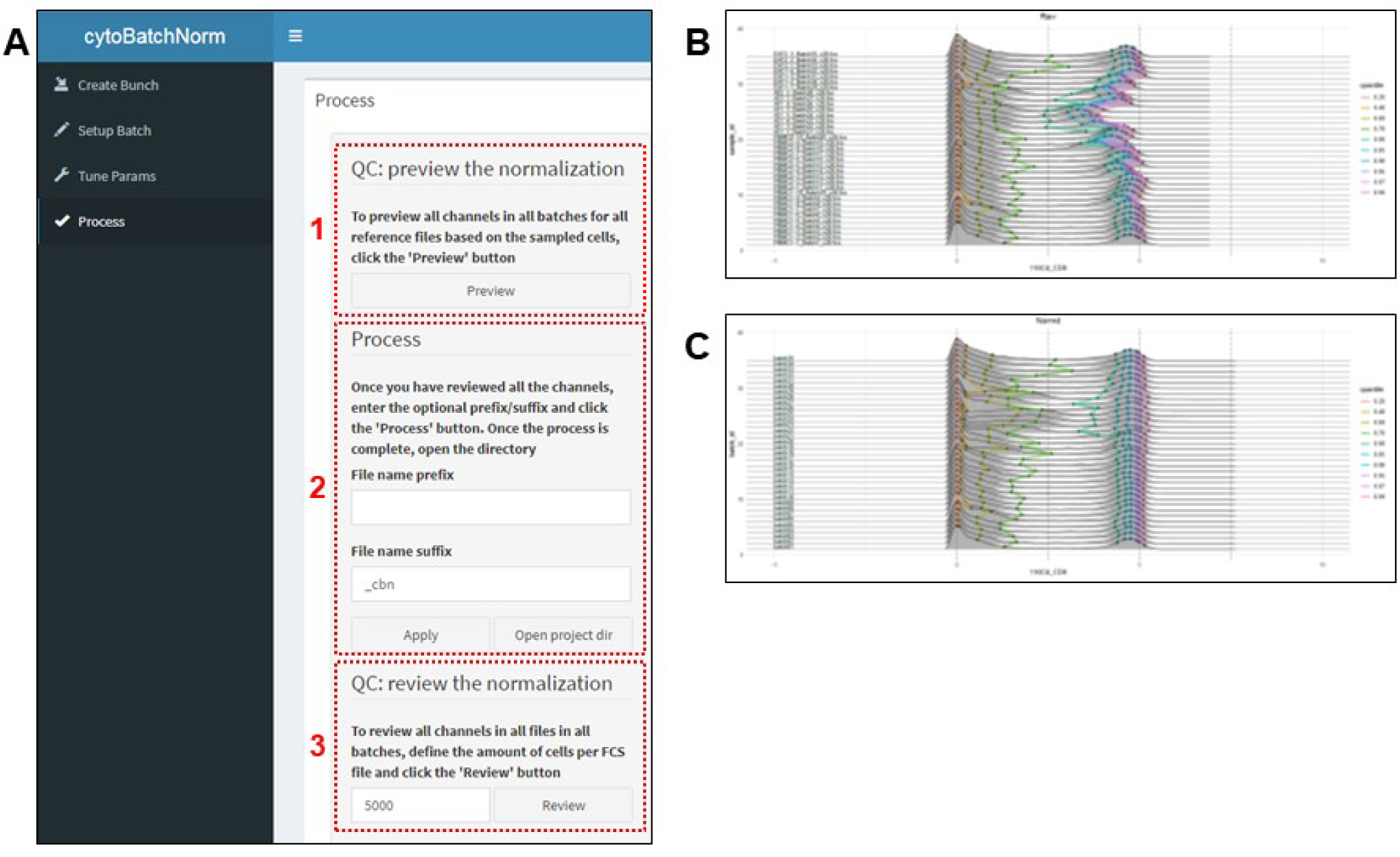
Detailed “process” tab. The Process menu allows to preview (A-1) the adjustment on all channels and all batches for reference files on PDF (B) consequently to parameters tuned previously. When satisfying, processing of FCS files for batch effects adjustments is launched by clicking on the “apply” button (A-2). Output channels adjustments can be reviewed on PDF files edited with le “review” button (A-3).

### “Process” menu

In the “Process” function (Figure 5), clicking on the “preview” button (Figure 5A-1) generates pdf files summarizing both raw (Figure 5B) and a preview of the adjustments to be realized on all control tubes (Figure 5C). A “prefix” and a “suffix” can be conveniently added to the output file names (Figure 5A-2). After having reviewed the normalization of the control samples, the normalization of all the FCS files is launched by clicking the “apply” button (Figure 5A-2). After calculation, the “review” button (Figure 5A-3) creates a pdf report summarizing channel histograms before and after adjustment (as a mirror of the “Tune parameters” menu) for each tube (control and experimental samples) of each batch, allowing a rapid visual control of the adjusted FCS files. Even if for some makers, this visual control is easily performed on histograms (as for CD8), checking adjusted FCS files (control tubes and experimental samples) on adequate and biologically-relevant dual plots is the best way to evaluate accuracy of the batch adjustment. Moreover, a quick manual gating (specifically for control of cytokines channels adjustment) is recommended. A summary of the final adjustments is exposed in Figure 6 (A and B), as well as downstream compensations (C) for Dataset 1.

**Figure 6:**
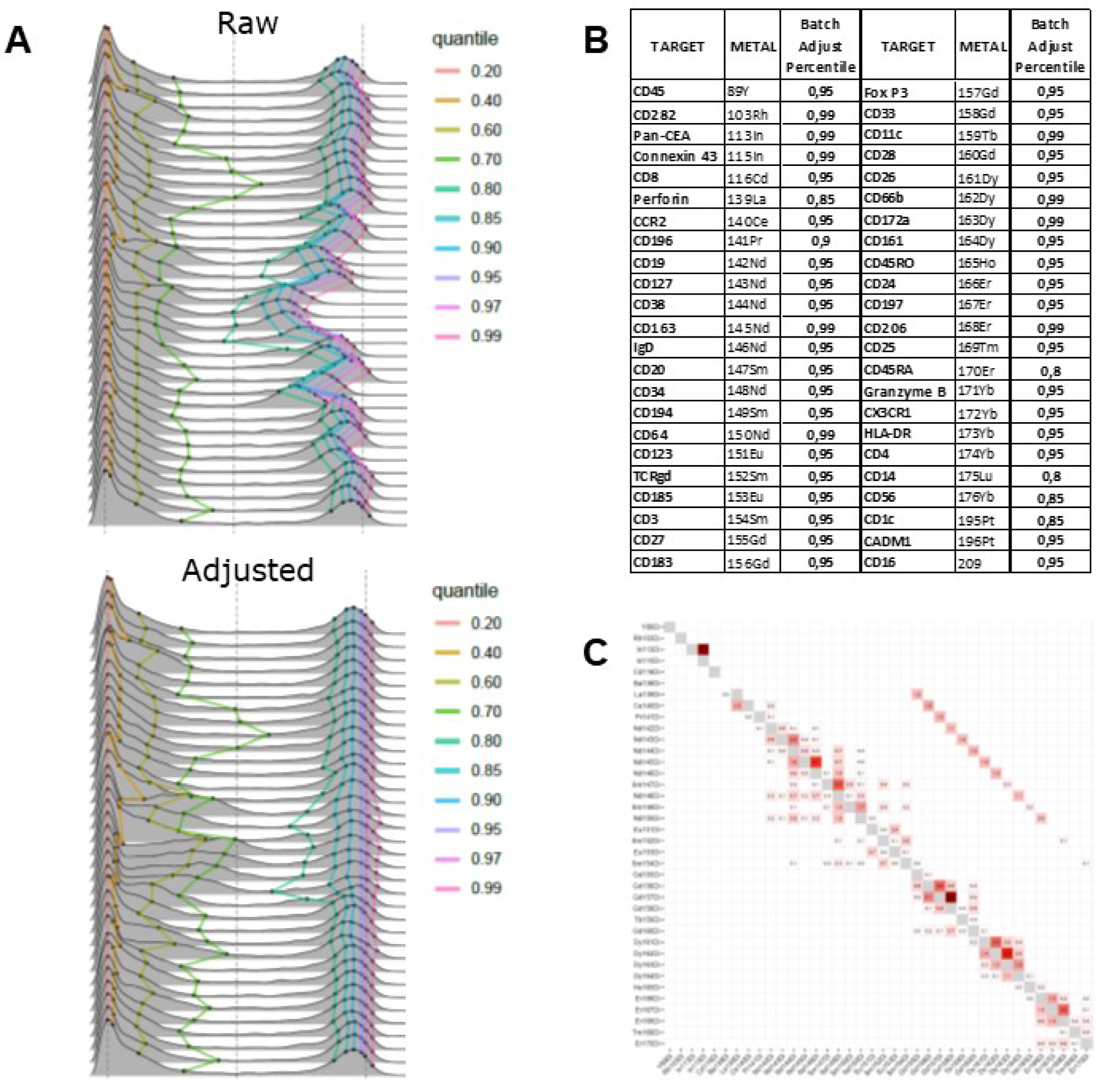
Mass cytometry data pre-processing. A: Illustration of batch adjustment on CD8 marker plots. B: Summary of percentiles used for batch adjustment of each channel of the panel. C: Compensation matrix calculated using CATALYST.

### Batch effects correction

Three levels of batch effects were identified and corrected in Dataset 1 as illustrated in Figure 7, termed as “single”, “barcode” and “time” batch effects. “Single” effects reflect specific normalization to a single batch for a given channel without any link to the batch or tissue-series it belongs to, as illustrated in Figure 7A (pink arrows). “Barcode” effects refer to batch homogenous specificity in correction levels for a given channel (Figure 7A, red brackets). The “time” effect was seen specifically in one experiment on PBMC (20 samples in one barcode), which displayed high variation of certain channels intensity during the 10 batch/runs of this barcode acquisition, as illustrated for CD45-_89_Y (Figure 7B). This probably reflects tolerable instability of the plasma torch, argon pressure, or TOF detector that were not corrected by normalization beads. These variations that were not corrected by the normalization on Four-Elements EQbeads (Standard BioTools Inc, San Francisco, CA, US) illustrates a very powerful implementation of batch correction for minimizing an instrument-related batch effect over time. Lastly, comparison of raw and batch-corrected control files on dimension reduction maps recapitulates the improvement of data homogeneity after processing to batch correction. Figure 7C illustrates accurate correction of batch effects-related CD8+ cells aggregation and expression level seen on a t-SNE map in the control sample from batch 23 (AT tissue, “Raw” vs “Normalized”) when compared to the reference batch 1. Multi-Dimensional Scaling (Figure 7D) using the CytoMDS R package[28] also illustrates reduced dispersion of AT samples (red circle) after correction (pink dots) compared to raw data (blue dots).

**Figure 7:**
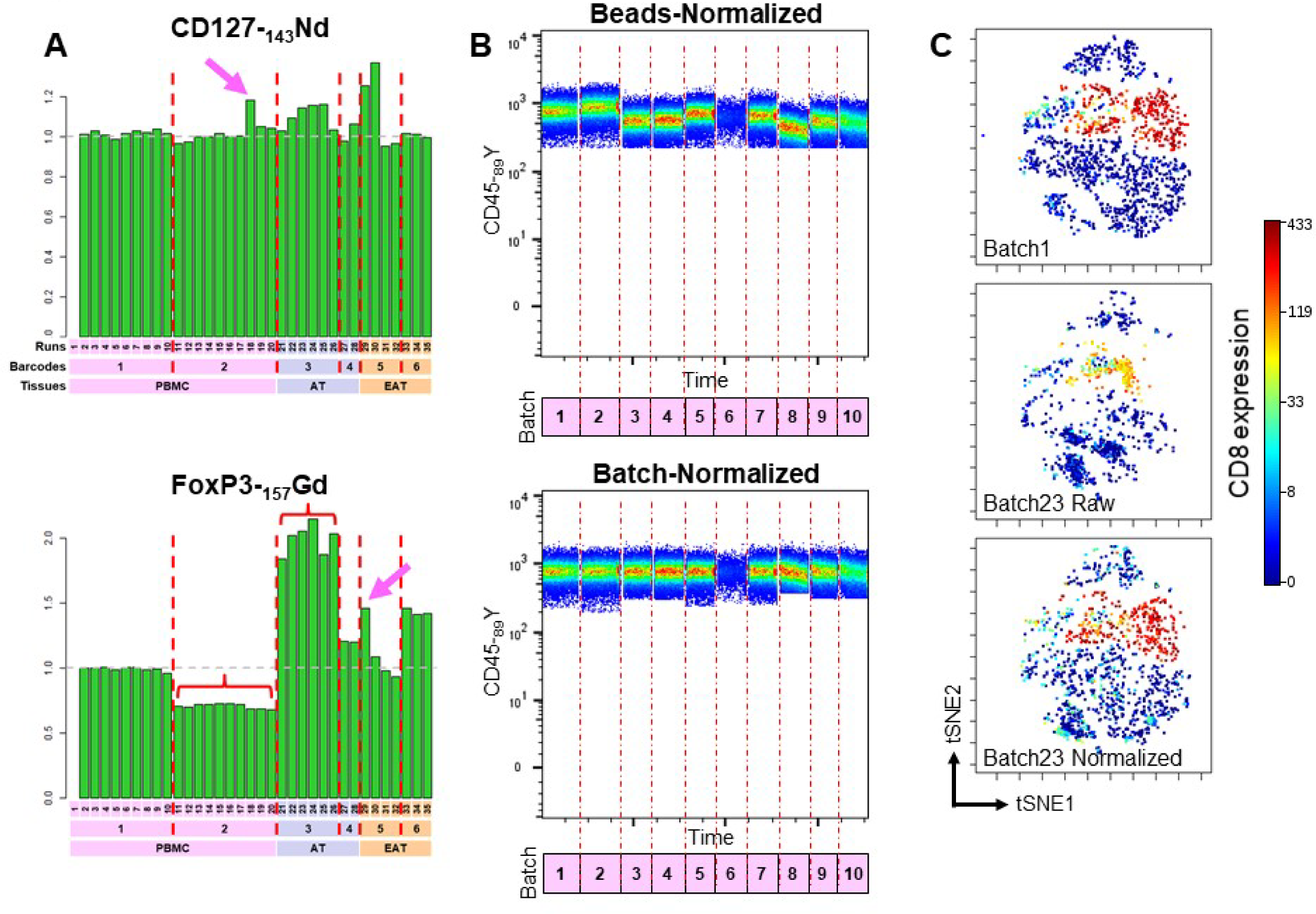
Different levels of corrected batch effects. A: scaling factors for CD127 and FoxP3 channels. Pink arrows illustrate “tube” specific batch effects; red brackets illustrate “barecode” specific batch effects. B: dot plots from PBMC batches 11 to 20 showing correction by CytoBatchNorm of basal CD45 levels variations over time (bottom plot) which were not corrected by beads normalization (top plot), illustrating “time” specific batch effects. C: CD8 expression on t-SNE dimension reduction map of lineage markers from control samples from Batch1 (PMBC) and Batch 23 (AT tissue) illustrates amelioration of data consistency following batch effect correction with CytoBatchNorm. D: Multi-Dimensional Scaling using the CytoMDS R package shows reduction of AT samples dispersion after batch effect correction (pink dots) compared to raw data (blue dots).

Finally, we adapted package code to allow its utilization on Windows systems.

### Benchmarking

Benchmarking of CytoBatchNorm was realized on Datasets 1 and 2 by two different scientists. Dataset 1 served for comparison between CytoBatchNorm and CytofBatchAdjust[19], while Dataset 2 served for comparison between CytoBatchNorm and CytoNorm[18].

As illustrated in Figure 8A, configuration handling time of CytoBatchNorm on a first attempt was about one hour for both datasets, reflecting time spent for reviewing all channels and configuring all percentiles of the dataset. On a second attempt, handling time decreases rapidly from one hour to 10-20 minutes because experimenters get used to both the interface and their dataset. This handling time on Dataset 2 was then twice that of CytoNorm, which methodology does not permit any tuning of channels corrections. CytofBatchAdjust correction to achieve the same level of fine tuning of channels adjustment (i.e. best individual channel percentile determination) requested numerous successive runs with different percentiles plus a systemic visual reviewing of the 700 FCS files composing Dataset 1 to determine the best percentile to choose, which took 16 hours. Whatever the package benchmarked, CytoBatchNorm calculation performed faster.

**Figure 8:**
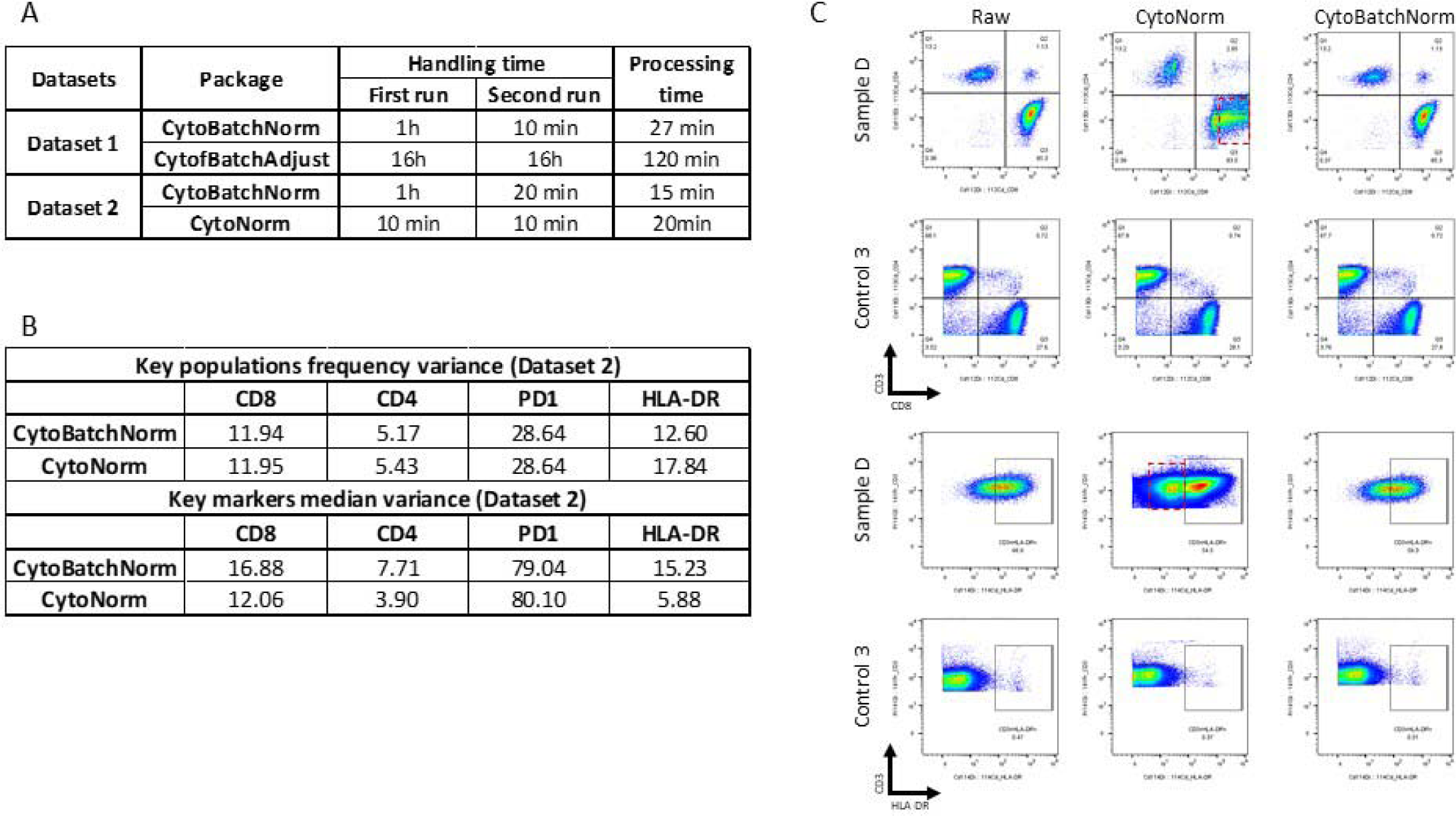
Benchmarking of CytoBatchNorm versus CytoNorm and CytofBatchAdjust. A: comparison of handling and processing time. B: Variance of key populations frequency and key markers median. C: Illustration of artifacts introduced in experimental samples by CytoNorm (red boxes).

Key populations variance comparison between Dataset 2 control files batch-corrected with either CytoBatchNorm or CytoNorm demonstrates CytoBatchNorm is at least as accurate as (CD8, PD1) or more accurate than (CD4, HLA-DR) CytoNorm (Figure 8B). On the opposite, CytoBatchNorm was slightly less accurate in reducing median variance of key markers (Figure 8B), except for PD1.

Strikingly, CytoNorm introduced artifactual deformation of some populations in some experimental samples, as illustrated in Figure 8C for CD8 and HLA-DR in sample 8 (red boxes) from the third batch of Dataset 2, but not in the batch specific control sample (CTL3). This illustrates 1) the absolute need for cautious control of potential bias introduced by specific algorithms miscomputation during data processing, 2) the inaccuracy of quantile-based normalization for cytometry data.

## CONCLUSION

Computer assistance in the treatment and analysis of omics data is ineluctable and has to be performed properly and wisely. Refining algorithms and free-access R packages to this aim will greatly enhance the still recent implementation of computational cytometry as well as the downstream results accuracy. One first, basic but essential step in cytometry data analysis is their standardization, including batch effects correction. We present the CytoBatchNorm R package which is the most user-friendly package available for batch effects correction, with a live assessment of correction accuracy, and which out-performs existing packages in terms of both tuning possibilities and efficiency. CytoBatchNorm will help the cytometry community to adequately scale their data amongst batches, allowing reliable reduction of variability and improvement of subsequent dimension reduction and clustering in user’s analysis pipeline.

## Abbreviations

AT: Atrial Tissue
CyTOF: Cytometry by Time Of Flight
EAT: Epicardial Adipose Tissue
PBMC: Peripheral Blood Mononuclear Cells
QC: quality control

## Acknowledgements

We thank Philippe Hauchamps (PhD student) from Computational Biology and Bioinformatics department, Duve Institute, Catholic University of Louvain, Belgium, for his active improvement of the CytoMDS R package used for figure 7D edition.

## Data Availability Statement

Due to confidentiality agreements, supporting data can only be made available to bona fide researchers subject to a non-disclosure agreement upon reasonable request. Details of the data and how to request access are available from at Univ. Lille, INSERM, CHU Lille, Pasteur Institute of Lille, U1011-EGID, Lille, France for Dataset 1, and from at Ludwig Institute for Cancer Research, Lausanne Branch, Department of Oncology, University of Lausanne (UNIL) and Lausanne University Hospital (CHUV), Agora Cancer Research Center, Lausanne, Switzerland for Dataset 2.

<u>Data availability statement</u>: as data serve only as an illustration dataset and are not yet published for scientific analysis of content, they are not made available nor submitted yet to flow repository.

## Ethics approval statement

The clinical cohorts were constituted as part of the POMI-AF study (NCT#03376165), approved by the institutional ethics committee (Comité de Protection des Personnes Ile de France V).

## Patient consent statement

Written informed consent was obtained from all patients before inclusion.

## Permission to reproduce material from other sources

None

## Clinical trial registration

None

