## Supplementary material for "CytoBatchNorm: an R package with graphical interface for batch effects correction of cytometry data": Figure S1

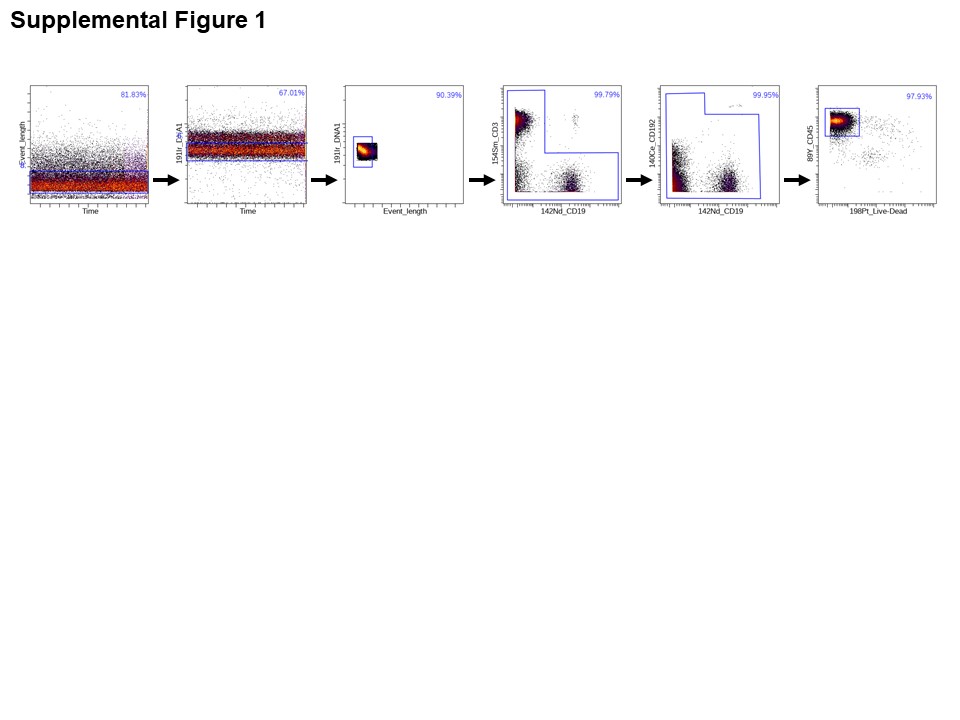

**Figure S1: Pretreatment of reference FCS files.** Manual gating of CD45^+^ live cells after exclusion of doublets, non-biological events and beads residues.
